# Effects of small-scale clustering of flowers on pollinator foraging behaviour and flower visitation rate

**DOI:** 10.1101/126581

**Authors:** Asma Akter, Paolo Biella, Jan Klecka

**Affiliations:** Biology Centre of the Czech Academy of Sciences, Institute of Entomology, České Budějovice, Czech Republic; Department of Zoology, Faculty of Science, University of South Bohemia, České Budějovice, Czech Republic

## Abstract

Plants often grow in clusters of various sizes and have a variable number of flowers per inflorescence. This small-scale spatial clustering affects insect foraging strategies and plant reproductive success. In our study, we aimed to determine how visitation rate and foraging behaviour of pollinators depend on the number of flowers per plant and on the size of clusters of multiple plants using *Dracocephalum moldavica* (Lamiaceae) as a target species. We measured flower visitation rate by observations of insects visiting single plants and clusters of plants with different numbers of flowers. Detailed data on foraging behaviour within clusters of different sizes were gathered for honeybees, *Apis mellifera*, the most abundant visitor of *Dracocephalum* in the experiments. We found that the total number of flower visitors increased with the increasing number of flowers on individual plants and in larger clusters, but less then proportionally. Although individual honeybees visited more flowers in larger clusters, they visited a smaller proportion of flowers, as has been previously observed. Consequently, visitation rate per flower and unit time peaked in clusters with an intermediate number of flowers. These patterns do not conform to expectations based on optimal foraging theory and the ideal free distribution model. We attribute this discrepancy to incomplete information about the distribution of resources. Detailed observations and video recordings of individual honeybees also showed that the number of flowers had no effect on handling time of flowers by honeybees. We evaluated the implications of these patterns for insect foraging biology and plant reproduction.

## Introduction

Plants typically vary in the number of flowers they produce and individuals often cluster together at various spatial scales. Clustered spatial distribution of flowers has implications both for plant reproduction and food intake of flower-visiting insects [1, 2]. Pollinator responses towards clustering of flowers at various spatial scales have long been studied and the outcomes are highly diverse. However, behaviour of flower visitors in relation to the number of flowers on individual plants, as well as their foraging behaviour in larger clusters of multiple plant individuals can be understood in the context of selection for behaviours maximising the efficiency of resource acquisition [3, 4]. Behavioural responses of pollinators to resource clustering at different spatial scales in turn affect reproductive success of plants [5–7].

At the scale of individual plants, pollinators often prefer to visit plants with a larger number of flowers. These provide a higher total amount of rewards (nectar and pollen); moreover, they can be detected from a larger distance [8]. Despite that, the number of visitors usually increases less than proportionally with the number of flowers [9–18]. Although pollinators generally visit more flowers in larger inflorescences, they tend to visit a smaller proportion of available flowers [10, 12–15, 17–19]. This behaviour is consistent with classic predictions of the optimal foraging theory, which assumes that foragers feed in such a way as to maximise their rate of net energy intake [3, 4]. When foraging in patches, they should leave after the rate of energy intake drops below the average level provided by other patches [20]. Because insects have a limited ability to remember which flowers they have already visited, they start to revisit empty flowers after some time [21]. The risk of revisiting empty flowers increases with increasing number of flowers per inflorescence [17, 22]. As a result, visiting a decreasing proportion of flowers in larger inflorescences is an optimal foraging strategy [21].

Each individual flower can be thought of as a small patch of food [10], where extracting nectar may become more difficult as the nectar is depleted. This could prompt the bee to move to another flower earlier in rich habitats to maximise the amount of food extracted per unit time [20]. Many invertebrates [23–25] and vertebrates [26, 27] feeding on various food sources were observed to shorten their handling time and discard partially consumed food items when food was abundant. However, this behaviour was not observed in previous studies on bees and syrphid flies [9, 22]. This suggests that these flower visiting insects may handle individual flowers in a constant manner independently of flower abundance, but more data are needed before drawing firm conclusions.

From the plant’s perspective, higher per-flower visitation rate should translate into higher reproductive output [28]. Most published studies found no relationship between the number of flowers in an inflorescence and per-flower visitation rate [14, 15, 17–19], although some reported an increasing [28] or decreasing [12] relationship. Moreover, the link between visitation rate and seed set is not straightforward. Percentage seed set may increase with the number of flowers when visitation rate also increases [16, 28], but it may be reduced in self-incompatible species due to geitonogamous pollination which occurs when a single pollinator visits multiple flowers on the same plant [15, 29, 30].

At the local scale, plants often grow in groups of multiple individuals, which we refer to as clusters. As in single plants, higher number of flowers in a cluster usually leads to a less than proportional increase in the number of visitors [10, 31–33], although proportional or higher increase was also reported [34]. Pollinators also tend to visit a decreasing proportion of flowers in larger clusters [10, 22, 31]. Visitation rate per flower usually stays constant because the increase in the number of visitors is counterbalanced by a decrease in the proportion of flowers visited by individual insects. This leads to the ideal free distribution of flower visitors [10, 17, 31, 33, 35]. At this spatial scale, optimal foraging theory is equally applicable for understanding flower visitor behaviour as in the case of individual plants with different sizes of inflorescences described above, and these patterns fit well to its predictions [10, 20–22]. However, the consequences for plant reproduction can be very nuanced. Percentage seed set was reported to be independent of cluster size [35], or increasing in response to higher visitation rate per flower in clusters with more flowers [34, 36]. However, seed set may also depend on the density of plants within the cluster [37], on their genetic compatibility [38], and on species-specific consequences of geitonogamous pollination whose frequency may vary with cluster size [30].

Here we report results of a field experiment conducted to test how flower visitation and foraging behaviour of pollinators depend on the number of flowers at two spatial scales: single plants and clusters of multiple plants. We conducted the experiment with potted *Dracocephalum moldavica* L. (Lamiacea). Specifically, we tested whether the number of visitors increases proportionally to the number of flowers on a single plant or in a cluster and whether plants with larger inflorescences or in larger clusters enjoy higher flower visitation rates. We then studied foraging behaviour of the most abundant flower visitor, *Apis mellifera*, in more detail to test how visit duration, number of flowers visited, and handling time per flower depend on the number of flowers. Our data show that the number of insects increased less than proportionally with the number of flowers and that honeybees visited a smaller proportion of flowers in larger clusters. Together, this led to maximal visitation rate per flower in clusters of intermediate size.

## Materials and Methods

*Dracocephalum moldavica* is a plant of the family Lamiaceae native to temperate zone of Asia; China, Russia, Tajikistan, and Turkmenistan. It is partly naturalised in a large part of Eurasia, introduced to the USA, and sometimes grown as an ornamental plant. It produces hermaphrodite flowers with violet colour which are oriented in whorls with 5-6 flowers in each whorl, have a semi-long corolla tube with nectaries at the bottom typical for Lamiaceae. The flower has a two-lobed stigma positioned below the upper lip and four anthers slightly shorter than the stigma. Each flower can produce four seeds. Interactions with pollinators are not well known; a related species, *Dracocephalum ryushiana*, is pollinated probably mostly by bumblebees [39]. We sowed the seeds in the beginning of May to germination trays in the greenhouse. Seedlings were transplanted individually to 1 litre pots containing a mixture of compost and sand (2:1) and grown in the greenhouse with daily watering. The plants fully flowered at the end of July with an average plant height of ca. 60 cm.

The first experiment (see below) was conducted in a meadow near Český Krumlov, 18 km southwest of České Budějovice (N 48°49.48’, E 14°18.98’). The rest of the project was carried out in a meadow near the campus of the University of South Bohemia in České Budějovice (N 48°58.50’, E 14°26.15’). All experiments were carried out on sunny days with no strong wind and no rain. No permits were required for this project because no protected species were collected and the study was conducted on public land.

### Experimental setup

The first experiment was designed to study pollinator visitation on single plants with different numbers of flowers (data in S1 Table). We used plants grown individually in pots. We adjusted the number of flowers per plant by cutting some of them, which provided plants with the number of flowers ranging from 1 to 174. Eight plants in pots were placed along a 35 m long transect; i.e. five meters apart. We observed and captured all flower visitors for 30 minutes per plant. Two to three people were collecting data simultaneously, each observing a different plant. We then replaced the plants by a new set of eight plants and repeated the observations. Overall, we sampled eight transects with different plant individuals during three days (4^th^, 7^th^, and 8^th^ August 2016), which resulted in a total of 64 observations. Sampling was conducted between 10:00 and 16:00 hours under good weather conditions (sunny, no rain). Insects were collected using an aspirator or a handnet, counted and preserved for identification.

The second experiment was aimed at studying visitation of clusters of multiple plants of different sizes (data in S2 Table). In this experiment, potted plants were placed to form five clusters 20 m apart in a 60 × 20 m grid (one position in the grid remained empty). Each cluster contained a different number of plants varying from 1 to 37. We also counted the number of open flowers at each plant. The number of flowers in a cluster ranged between 42 to 2476. Each cluster was observed for 30 minutes during which all insects visiting *Dracocephalum* flowers in the cluster were captured and preserved. We completed seven sampling periods on 16^th^ and 17^th^ August 2016, which yielded in total 35 observations of cluster visitation. The numbers of flowers in each cluster were counted every day after finishing the experiments.

We also conducted detailed observations of foraging behaviour of *Apis mellifera* at the site of the second experiment (data in S3 Table). The total number of flower visitors in the experiments was dominated by *Apis mellifera*, which was thus selected for additional measurements. We measured the duration of visits, the number of flowers exploited, and handling time per flower of *A. mellifera* in clusters of different sizes. Potted plants were placed in the same grid as in the second experiment to form five clusters on 25^th^ and 26^th^ August 2016. The number of plants per cluster ranged from 1 to 22, and the number of flowers was 2 to 643 per cluster. In these observations, a single *A. mellifera* was followed from its entry into the cluster until its last visit to a flower in the same cluster. Data collection included both the time spent in one cluster measured by a stopwatch and the number of flowers visited by each individual *A. mellifera*. To test the hypothesis of partial consumption in honeybees, we measured the average number of flowers per minute over individual foraging bouts based on direct observations and then measured time spent on individual flowers using video recordings (data in S4 Table). Video recordings were taken at the same time as observations of honeybee foraging, but in those clusters which were not observed at the time to minimise disturbance of the recordings. We distinguished: i) total time from landing at a flower until leaving and ii) actual feeding time (head inserted deep inside the flower). We tested whether these two measures of handling time depended on the number of flowers in a cluster.

### Data analyses

For the experiments on flower visitation, we conducted the analyses at the level of the total number of insect visitors per plant or per cluster. We tested how the number of flower visitors and other measures of visitation varied with the number of flowers using generalised linear models (GLM), or generalised additive models (GAM) implemented in a package mgcv 1.8-17 [40] when the relationship was nonlinear. Analyses were done in R 3.2.4 [41]. The number of flowers, used as an explanatory variable, was log-transformed before the analyses. We fitted the GLMs and GAMs using overdispersed Poisson distribution (quasipoisson) or Gamma distribution with log link function depending on the response variable. Analysis of proprotion data was performed using Beta regression implemented in betareg package for R [42].

## Results

We found that the number of flower visitors increased with the increasing number of flowers on a plant or in a cluster, but less than proportionally (GLM, quassipoison distribution, *F*_1,62_ = 31.5, *P* < 10^−6^; Fig 1A, blue line and points). The relationship was linear at the log-log scale with a slope of 0.57 (*SE* = 0.108). Data from larger clusters of multiple plants qualitatively showed an extension of the patterns observed in single plants. The number of insects increased with the increasing number of flowers (*F*_1,33_ = 45.6, *P* < 10^−6^; Fig 1A, orange line and points) with a slope of 0.58 (*SE* = 0.093) at the log-log scale.

**Fig 1.**
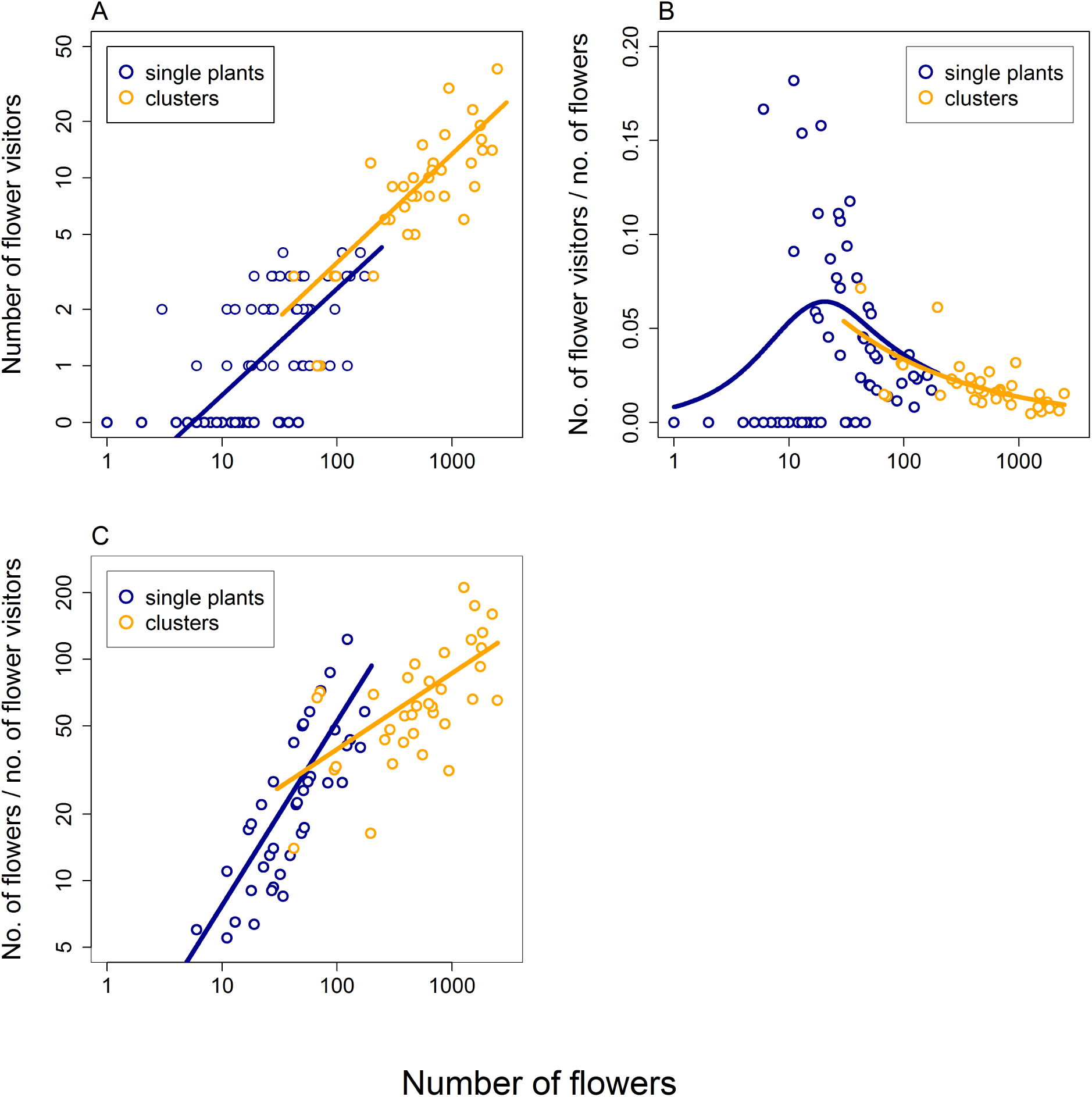
The effects of the number of flowers in single plants and larger clusters on visitation by insects. The plots combine data from two separately conducted experiments: one with single plants differing in the number of flowers (blue circles and fitted lines) and another with larger clusters of up to 36 plants (orange circles and fitted lines). Data from these two experiments were combined for the purpose of visualisation, but were analysed separately. A: The number of flower visitors observed during 30 minute observation periods on single plants and larger clusters varying in the number of flowers. B: The number of insects visiting the plants or clusters of multiple plants relative to the number of flowers available. C: The number of flowers available per visitor; i.e. the potential pay-off for the flower visitors.

Also, the ratio between the number of visitors and flowers increased when the number of flowers increased from one to around 20, but decreased when the number of flowers increased further (GAM, Gamma distribution, log link function, *F* = 3.2, *P* = 0.0026; Fig 1B, blue line and points). The log-linearly decreasing relationship in clusters of plants was an extension of the relationship reported in single plants (*F* = 26.1, *P* = 1 × 10^−5^; Fig 1B, orange lines and points).

We also calculated potential payoff for flower visitors defined as the mean number of flowers per visitor, assuming that already visited flowers did not renew their nectar reward during the observation time (30 minutes). The potential payoff increased with the increasing number of flowers on both single plants (GLM, Gamma distribution, log link function, *F*_1,39_ = 67.5, *P <* 10^−6^; Fig 1C, blue line and points) and clusters of plants (*F*_1,33_ = 21.0, *P* = 6 × 10^−5^; Fig 1C, orange line and points), although with a shallower slope in the latter case; slope was 0.83 in single plants (*SE* = 0.096) and 0.34 in clusters of plants (*SE* = 0.076). This means that there were more free resources available for each visitor in larger clusters.

Detailed observations of foraging behaviour of individually tracked honeybees showed that, as expected, individual honeybees spent more time foraging in larger clusters (GLM, Gamma distribution, log link function, *F*_1,80_ = 8.5, *P* = 0.0045; Fig 2A) and visited more flowers there (GLM, quasipoisson distribution, *F*_1,80_ = 11.3, *P* = 0.0012; Fig 2B). However, the increase was only modest in both cases; significantly less than proportional. The slope was 0.34 (*SE* = 0.114) for time and 0.38 (*SE* = 0.117) for the number of flowers visited. There was also considerable variation around the fitted relationships. The proportion of available flowers visited by individual honeybees decreased significantly with the increasing number of flowers per cluster (Beta regression, *χ*^2^ = 8.9, *P* = 0.0029; Fig 2C). In large clusters, all individuals visited only a minority of flowers, while in small clusters, the proportion of flowers visited varied widely from just a few to all flowers available (Fig 2C). Honeybees foraged with the same speed across the range of cluster sizes; i.e. the number of flowers visited per minute did not depend on the number of flowers available (GLM, Gamma distribution, log link function, *F*_1,80_ = 1.87, *P* = 0.1756, Fig 2D).

**Fig 2.**
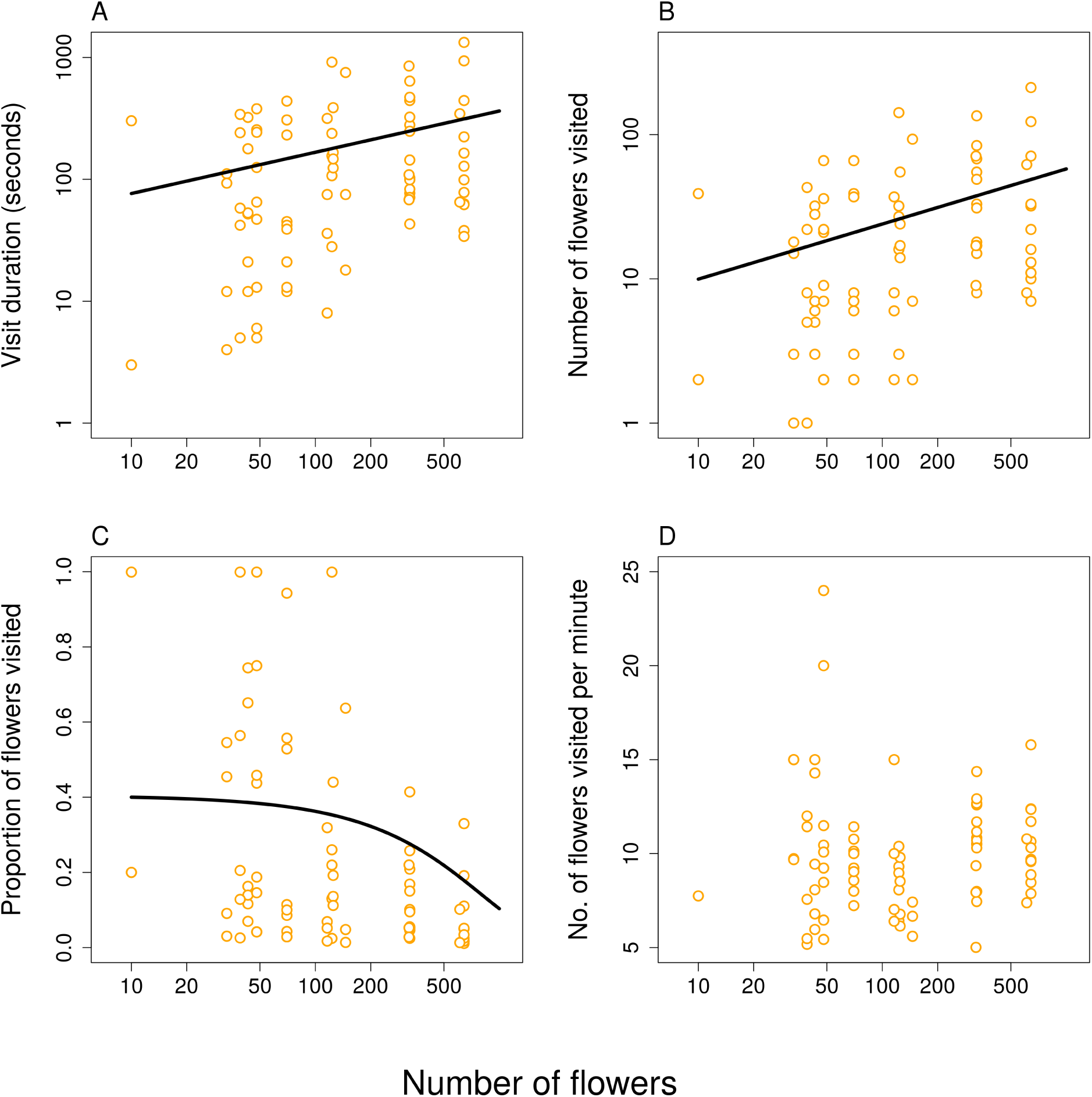
Foraging behaviour of honeybees in response to the number of flowers in clusters of multiple plants. A: The time spent foraging within the cluster by individual honeybees increased with the number of flowers available. B: The number of flowers visited increased with the total number of flowers available. C: The proportion of flowers visited by individual bees decreased with the number of flowers available. D: The number of flowers exploited per minute did not show any significant relationship to the number of flowers available. X-axis in A. - D. and y-axis in A. and B. are on a log-scale.

Neither of our two measures of handling time per flower depended on the number of flowers in a cluster (Fig 3). Feeding time per flower, defined as the time a bee spent with its head deep inside a flower, apparently engaged in nectar extraction, did not depend on the number of flowers in a cluster (GLM, Gamma distribution, log link function, *F*_1,132_ = 0.75, *P* = 0.3869; Fig 3A). Also the total time spent on the flower from first contact until take-off was independent of the number of flowers (GLM, Gamma distribution, log link function, *F*_1,132_ = 0.95, *P* = 0.3308; Fig 3B). Compared to data shown in Fig 2D, these measurements exclude travelling time between flowers and estimate only the time spent handling the flowers and the actual duration of feeding per flower.

**Fig 3.**
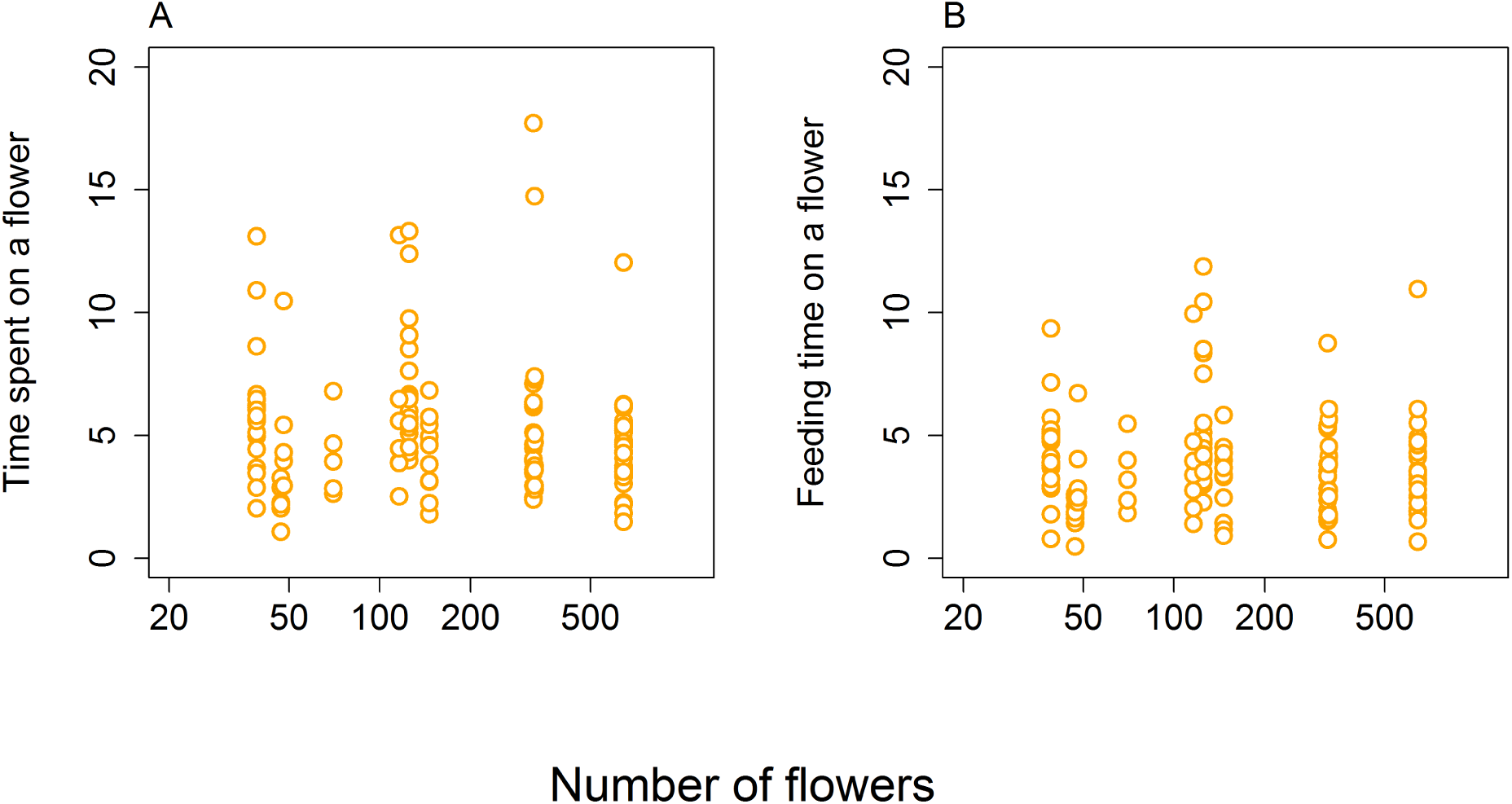
Honeybees’ handling time per flower was independent of the number of flowers in clusters of different sizes. A: Total time spent on a flower measured from video recordings as the time from the first contact until take-off. B: Feeding time estimated as the time honeybees spent with their head deep inside the flower. Both are measures of handling time excluding movement between flowers.

There was a considerable variation in both total time spent on a flower and feeding time per flower in individual flower visits, which was quantified by generalised linear mixed models. We fitted a model including the intercept and individual identity as a random factor with log-transformed response variables to quantify the variance of handling time between individuals and within individuals. Based on this, we calculated the ratio of between-individual variance and total variance, i.e. repeatability, as a measure of differences between individuals (rptGaussian function in rptR package for R, [43]). We found that repeatability of the total time spent on a flower was 0.15 (95% confidence interval = [0, 0.321] based on bootstrap); i.e. 15% of the variance occurred at the between-individual level and 85% at the within-individual level. Repeatability of the feeding time was only slightly higher, 0.21 (95% confidence interval = [0.0346, 0.405] based on bootstrap). There were thus only small differences between individuals in both measures of their handling times.

Finally, we calculated visitation rate per flower and unit time by multiplying the estimated dependence of the number of honeybees on the number of flowers (GLM, quassipoison distribution, *F*_1,33_ = 10.009, *P* = 0.0034, slope=0.40, *SE* = 0.1313) and the dependence of the proportion of flowers visited on the number of flowers per cluster (Fig 2C). The estimated visitation rate showed a unimodal relationship peaking at the intermediate level of the number of flowers per cluster (Fig 4).

**Fig 4.**
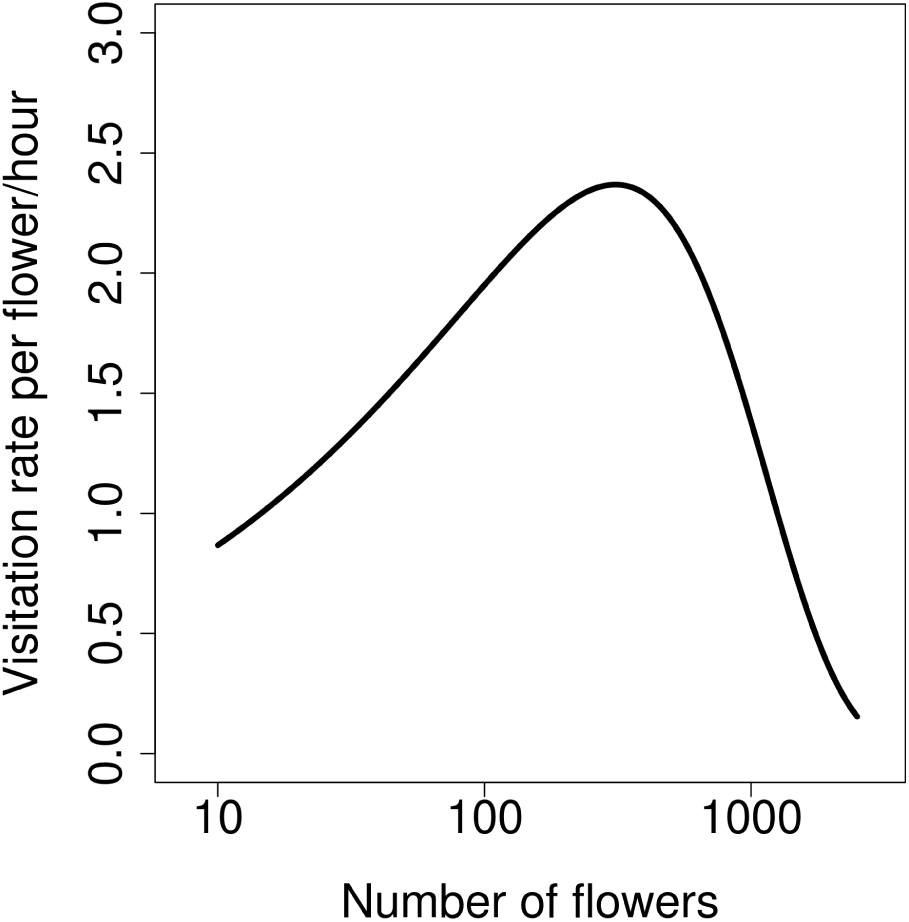
Flower visitation rate peaks at the intermediate number of flowers. Visitation rate per flower per hour was estimated as a product of the number of visitors (honeybees only) and the proportion of flowers visited by an individual honeybee.

## Discussion

We observed that single plants with many flowers and large clusters were generally more attractive to flower-visiting insects than those with a smaller number of flowers. This is a classic pattern expected for optimally foraging animals who maximise net energy intake per unit time. However, our data show several departures from simple theoretical expectations.

The number of flower visitors increased with the increasing number of flowers, but less than proportionally (Fig 1A). Optimally foraging animals should reach *ideal free distribution* (IFD) where they would possibly ignore very poor patches altogether, and they would be distributed between the rest of the patches in such a way as to equalise patch payoff [10, 44]. In the case of flower visitors, this leads to flower visitation rate independent of the number of flowers per plant or cluster [10]. However, our data show that plants with many flowers and large clusters were underutilised. The number of insects per flower decreased sharply in single plants with many flowers and in large clusters (Fig 1B), and the number of flowers available per visitor increased (Fig 1C). Detailed observations of foraging honeybees, the most numerous flower visitor species, showed that flower visitation rate peaked in clusters of plants with an intermediate number of flowers and dropped in clusters with both few flowers and many flowers (Fig 4). This observation is inconsistent with the prediction of a constant flower visitation rate based on optimal foraging theory [10]. Previous empirical studies generally found that i) the number of visitors increased less than proportionally with the number of flowers and ii) that an increasing number of flowers was visited per individual in larger clusters. Our results also show these patterns. In a number of previous studies these two relationships had such slopes that they resulted in constant flower visitation rate [10, 17, 31, 33, 35]. However, in our case, these relationships had such shapes that they combined to form a unimodal pattern with the highest flower visitation rate in clusters with an intermediate number of flowers (Fig 4). This represents suboptimal foraging behaviour because large, most profitable clusters were underutilised. Other reported deviations from the expected pattern include a decreasing [12] as well as increasing [28] flower visitation rate in larger clusters.

The lack of flower-visitors on plants with few flowers is consistent with expectations based on optimal foraging theory [20] and the IFD model [44]. It is generally not profitable to use poor resources, i.e. plants with few flowers, unless resources are very scarce [44]. Honeybees are known to adjust their selectivity for clusters of flowers based on the overall abundance of resources, so they avoid poor resources when food is plentiful [45]. However, an alternative explanation is that this is not due to choice on the part of insects but due to low detection probability of plants or clusters with few flowers. Detectability of an object increases with the visual angle subtended by the stimulus, which means that bees and other animals can see large flowers or inflorescences easier and from a larger distance [8, 46, 47]. Unfortunately, our data do not allow us to decide whether plants with few flowers were not detected or ignored.

Underutilisation of plants and clusters with a high number of flowers could be explained by a limited amount of information insects had about the quantity and spatial distribution of resources, because we placed the plants at the meadow only shortly before we started our observations [48–51]. Classic models of optimal foraging theory [20] and IFD [44] assume that foragers are omniscient, i.e. that they know the quality of all individual patches of food. This is rarely if ever the case in reality, so animals must make foraging decisions with imperfect information [48–50, 52]. They are generally thought to use information about the quality of previously visited clusters together with their perception of the quality of a new cluster to decide whether to enter the cluster or go elsewhere [48, 50]. This may provide explanation for our observation of underutilisation of the richest clusters. The meadow where the experiment was conducted had fairly low abundance of flowers, so medium-sized and larger clusters of *Dracocephalum* probably provided a richer source of food than the original plant community. At the same time, insects had a limited amount of information about the location and quality of the clusters of *Dracocephalum*. Our experimental manipulation thus represents a case of quick changes in resource availability and spatial distribution, similar to common natural situations such as when some plants start flowering and the spatial distribution of resources for flower-visiting insects changes over short time-scales. In such cases, bees have a limited amount of information about their resources, so they are not able to forage optimally [53]. Our data thus support previous observations that foragers are usually overrepresented in poor clusters and underrepresented in rich clusters, leading to suboptimal food intake [48].

At the within-cluster scale, we observed that individual honeybees spent more time and visited more flowers in larger clusters, but they visited a smaller proportion of the available flowers (Fig 2). This pattern has already attracted considerable attention because it seems to be at odds with optimal foraging behaviour [10, 22, 31, 54]. However, due to larger numbers of insects visiting larger clusters of flowers, visiting a smaller proportion of flowers leads to an IFD and thus to an optimal use of resources [10]. For example, Goulson [22] performed experiments which showed that as the insect visits flowers in a large patch it becomes difficult to avoid revisiting already emptied flowers, so at some point it becomes advantageous to leave the patch rather than search for the remaining unvisited flowers because food intake rate is depressed [20]. Another aspect of foraging biology we studied was handling time per flower. None of the measures we used varied with the number of flowers per cluster (Fig 2D and Fig 3), so it appears that bees did not adjust the way they used individual flowers depending on the number of flowers in a cluster. This result is in line with several previous studies on various bees and syrphid flies [9, 22], so these insects apparently handle individual flowers in a constant manner independently of flower abundance. It is important to note that most studies of foraging behaviour focused on honeybees or bumblebees, which may behave differently from other groups of pollinators. For example, it seems that honeybees visit a higher proportion of flowers before moving to another plant compared to other pollinators [55]. Comparative studies on multiple flower visitors will be needed to shed more light on the generality of patterns discussed here, see e.g. [9].

Our current data do not allow us to evaluate the implications of flower clustering for plant reproduction. Our observations of flower visitation and foraging behaviour suggest that the number of flowers on individual plants and in clusters of plants could affect plant reproductive success. Specifically, there was a lower number of insects per flower on plants with more flowers and in larger clusters (Fig 1) and per-flower visitation rate peaked in clusters with an intermediate number of flowers (Fig 4). Variation in flower visitation rate should lead to differences in pollination and consequently percentage seed set depending on the number of flowers per plant or per cluster. However, previous studies show that the link between visitation and seed set in plants is often weak and not at all straightforward [32, 56]. Additional data from a different type of experiments will thus be needed to resolve this question in our system.

## Conclusions

Our results show that flower-visiting insects preferred plants and clusters of multiple plants with larger numbers of flowers. However, visitation rate per flower and unit time peaked in clusters with an intermediate number of flowers in violation of ideal free distribution expected for optimally foraging animals. We consider imperfect information about the location and quality of plant clusters to be a likely explanation of this pattern. Detailed observations of foraging honeybees showed that they visited more flowers, but a smaller proportion of flowers in larger clusters. Finally, although handling time per flower was highly variable, it was unrelated to the number of flowers per cluster. Bees were thus not flexible in handling flowers depending on their local abundance.

## Supporting information

**S1 Table. Data from the first experiment on flower visitation of individual plants with a variable number of flowers**.

**S2 Table. Data from the second experiment on flower visitation of clusters of multiple plants**.

**S3 Table. Detailed data on foraging behaviour of honeybees**.

**S4 Table. Data on honeybee handling time per flower: feeding time and total time spent per flower**.

## Acknowledgements

We would like to thank Pavla Dudová for help with field observations. We are also grateful to two anonymous reviewers who provided valuable comments on the manuscript.

